# Inferring Spatially Resolved Transcriptomics Data from Whole Slide Images for the Assessment of Colorectal Tumor Metastasis: A Feasibility Study

**DOI:** 10.1101/2022.11.24.517856

**Authors:** Michael Fatemi, Eric Feng, Cyril Sharma, Zarif Azher, Tarushii Goel, Ojas Ramwala, Scott Palisoul, Rachael Barney, Laurent Perreard, Fred Kolling, Lucas A. Salas, Brock C. Christensen, Gregory Tsongalis, Louis Vaickus, Joshua J. Levy

## Abstract

Over 150,000 Americans are diagnosed with colorectal cancer (CRC) every year, and annually over 50,000 individuals will die from CRC, necessitating improvements in screening, prognostication, disease management, and therapeutic options. Tumor metastasis is the primary factor related to the risk of recurrence and mortality. Yet, screening for nodal and distant metastasis is costly, and invasive and incomplete resection may hamper adequate assessment. Signatures of the tumor-immune microenvironment (TIME) at the primary site can provide valuable insights into the aggressiveness of the tumor and the effectiveness of various treatment options. Spatially-resolved transcriptomics technologies offer an unprecedented characterization of TIME through high multiplexing, yet their scope is constrained by cost. Meanwhile, it has long been suspected that histological, cytological and macroarchitectural tissue characteristics correlate well with molecular information (e.g., gene expression). Thus, a method for predicting transcriptomics data through inference of RNA patterns from whole slide images (WSI) is a key step in studying metastasis at scale. In this work, we collected and preprocessed Visium spatial transcriptomics data (17,943 genes at up to 5,000 spots per patient sampled in a honeycomb pattern) from tissue across four stage-III matched colorectal cancer patients. We compare and prototype several convolutional, Transformer, and graph convolutional neural networks to predict spatial RNA patterns under the hypothesis that the transformer and graph-based approaches better capture relevant spatial tissue architecture. We further analyzed the model’s ability to recapitulate spatial autocorrelation statistics using SPARK and SpatialDE. Overall, results indicate that the transformer and graph-based approaches were unable to outperform the convolutional neural network architecture, though they exhibited optimal performance for relevant disease-associated genes. Initial findings suggest that different neural networks that operate on different scales are relevant for capturing distinct disease pathways (e.g., epithelial to mesenchymal transition). We add further evidence that deep learning models can accurately predict gene expression in whole slide images and comment on understudied factors which may increase its external applicability (e.g., tissue context). Our preliminary work will motivate further investigation of inference for molecular patterns from whole slide images as metastasis predictors and in other applications.

## 1. Introduction

Colorectal cancer is the third leading cause of cancer-related death in the United States, and there are disparities in screening and outcomes between age, race, and gender ^1^. CRC incidence is rising among younger adults who are not typically incorporated into screening programs, illustrating the importance of developing timely and lower-cost prognostication methods to better assess the tumor’s malignant potential. Currently, the Pathological TNM-staging system (pTNM), which determines staging based on the impact of local invasiveness–histological stage (T-stage), and metastatic potential– nodal (N-stage) and distant (M-stage) metastasis, is the most predictive factor for risk of recurrence and prognosis. Metastasis, in many cases, is challenging to assess at the time of primary tumor resection ^2^. For instance, specimen inadequacy often hinders the complete inference of nodal involvement ^3^. It is thus crucial to develop orthogonal, less invasive, but equally informative technologies that can shed light on disease pathology and prognosis. One promising direction is to study the Tumor Immune Microenvironment (TIME)– the amalgamation of immune cells, chemokines, cytokines, and other immune modulators, etc. that accrete at the invasive front and inside the tumor at the primary site ^4–6^. Recent studies have demonstrated that monocyte/lymphocyte immune infiltrates and their spatial distribution, density, and relationships play an important role in providing a coordinated anti-tumoral response. Yet, the full importance of TIME has not been elucidated, as most clinical studies consider either a few canonical markers at a time (e.g., immunoscore, which assesses cytotoxicity at the primary site) or only study cell mixtures which lack a single-cell or spatial dimension ^7^.

Spatially-resolved transcriptomics (spatial omics), as enabled through technologies such as 10x Genomics Spatial Transcriptomics (ST, Pleasanton, CA) or Nanostring GeoMX Digital Spatial Profiling (DSP, Seattle, WA), is an actively growing area of research that provides rich information about how different areas of tissue interact by analyzing highly multiplexed gene expression at staggering spatial resolution. These technologies can be configured to study the distribution, density, and spread of tumor-infiltrating lymphocytes (TILs) as they may relate to concomitant somatic alterations ^8,9^. Assay costs are currently exceedingly high, as profiling just four capture areas can cost tens of thousands of dollars, though costs are being driven downwards with new advances in chemistry and lower sequencing costs. Thus, sufficiently powering spatial transcriptomic association studies or extending their generalizability of the inferences to specific patient subgroups that lie outside of these small cohorts is challenging due to cost. In comparison, tissue slides stained with hematoxylin and eosin (H&E) to assess tissue histomorphology are routinely ordered at a very low cost, and there is ample evidence to suggest that many concurrent molecular alterations coincide with morphological features. Thus, the prediction of RNA expression using image data across a slide presents an opportunity to reveal critical prognostic information for patients at a lower cost, which can motivate relevant downstream analyses.

Deep learning approaches, which rely on using multi-layer artificial neural networks (ANN), have proven instrumental for image analyses in the context of digital pathology ^10^. Of relevance for this study is the assessment of whole slide images (WSI), digitized tissue slides, from which machine learning applications can predict the primary site of a metastatic lesion, tumor stage, and the outcome of immunohistochemical stains. Convolutional neural networks (CNNs), a type of predictive machine learning model, are powerful tools for extracting dense information from high-dimensional image data. Prior works have employed these algorithms to extract morphological features from H&E-stained tissue to complement whole transcriptome analyses. As WSI can extend to hundreds of thousands of pixels along each spatial dimension, they are usually broken into subimages to enable efficient computation. Of relevance to our research topic, He et al. (2020) used a DenseNet-101 model to regress on co-localized gene expression levels^11^, and Levy-Jurgensen et al. (2020) employed an InceptionV3 model to detect dichotomized gene expression for given patches of tissue^12^. However, these techniques do not analyze the potential for integrating spatial context outside the patch-level (i.e., spatially correlated patches are assumed to be independent and identically distributed), and, therefore may miss larger macroarchitectural contextual cues of aberrant expression.

Zeng et al. (2022) and Pang et al. (2021) investigated approaches to integrate broader spatial context into their gene expression prediction models by using vision transformers and demonstrated that it is possible to outperform convolutional architectures that make predictions on individual patches (e.g., ST-Net) ^13,14^. However, there remain several unknowns, such as 1) the scale of tissue features relevant for inferring RNA, 2) how well these models preserve global spatial expression characteristics (e.g., patterns of clustering), 3) whether relevant domain knowledge can make inferences more informative, and 4) whether these effects depend on the specific genes under study (i.e., what resolution/architectural context is optimal for specific genes and what does it say about the tumor biology). Furthermore, many spatial omics studies attempt to study one slide, where potential endogeneity is introduced by a matter of spatial distance between collection sites.

Here, we compared a convolutional neural network and a graph attention network for inferring spatially co-registered gene expression from WSI. We comment on the role the tissue macroarchitecture may play in RNA inference as elucidated by changing subimage size and the use of contextual models. We also assessed performance on a subset of immune-related genes and how spatial characterization varies across these models/factors and slides. These explorations will motivate future downstream work to characterize factors pertaining to tumor nodal/distant metastasis.

## 2. Material and Methods

### 2.1. Overview

The primary goal of this work is to predict the gene expression detected by a Visium spot at any given location on the slide. Our method is as follows (**Figure 1**):

1. **Data Collection:** Acquire H&E whole slide images (WSI), and spatially-registered Visium assayed spatial transcriptomics slides from 4 stage-pT3 matched colorectal cancer patients at Dartmouth Hitchcock Medical Center, two patients without metastasis, one with nodal metastasis only and one with both nodal and distant metastasis.
2. **Preprocess**: Preprocess gene expression and WSI subarrays.
3. **Model Development**: Configure two modeling approaches: convolutional neural network and graph attention neural networks, the latter leveraging larger spatial context. These models will predict both binary (dichotomized expression) and count-based (continuous; zero-inflated negative binomial) objectives. We also configured additional approaches (e.g., Transformer) for comparison.
4. **Capture surrounding tissue context at different scales**: Ablation study over patch size to determine whether relevant biological information is encoded outside the Visium collection spot area.
5. **Leave one-patient-out cross-validation**: Evaluation on held-out slides/patients as a measure of external applicability.
6. **Recover Spatial Biology Inferences:** Spatial autocorrelation tests for the capacity of models to draw similar spatial inferences.

**Figure 1.**
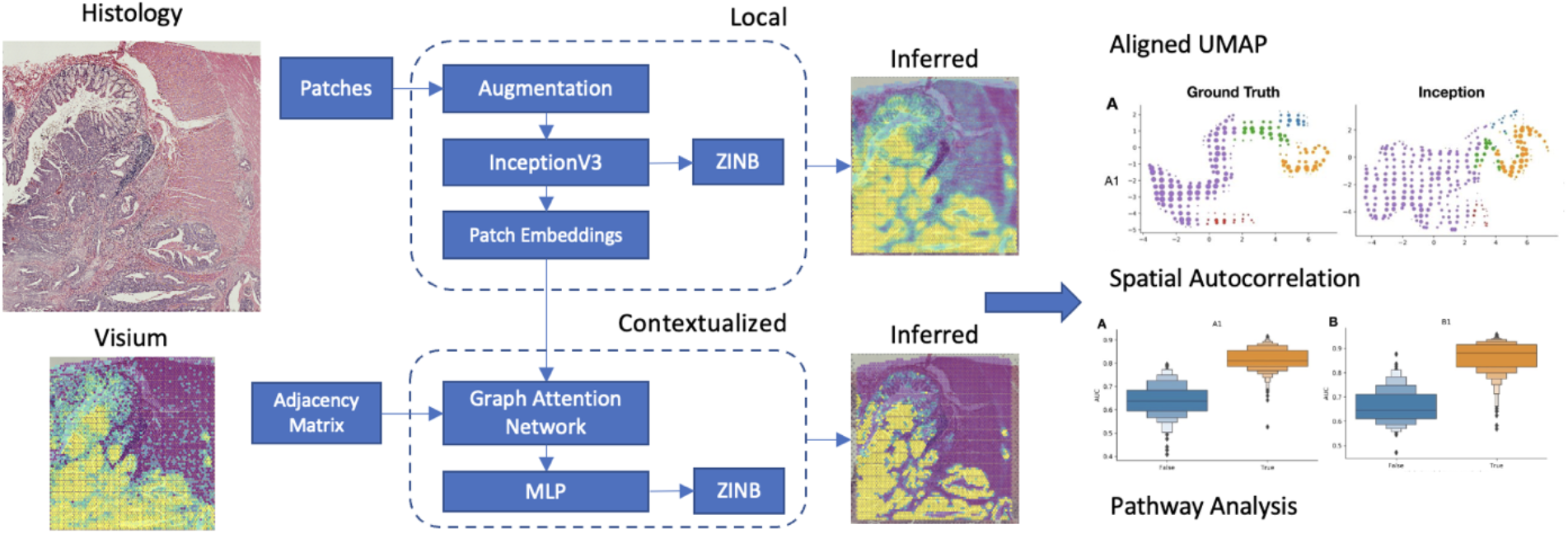
Overview of tested model architectures: Whole Slide Images are divided into patches co-localized with the Visium spots and an Inception model is used to predict counts and dichotomized expression for 1000 genes; Features derived from Inception model are additionally fine-tuned using graph neural network for inference; inferred expression profiles are compared to the ground truth through a cluster analysis, spatial autocorrelation tests and pathway analysis

### 2.2. Data Collection

The primary dataset utilized in this study was acquired from four pathologic T Stage-III (pT3) matched (pTNM system) colorectal cancer patients at Dartmouth Hitchcock Medical Center, determined through a retrospective review of pathology reports from 2016 to 2019 following IRB review and approval. These four patients were drawn from a set of 36 patients included in a prior study ^15^– half of the patients had concurrent tumor metastasis (slides A1 and B1 had tumor metastasis; slides C1 and D1 did not have tumor metastasis) and were otherwise matched on age, sex, tumor grade, tissue size, mismatch repair (MMR) status, and tumor site using iterative patient resampling with t-tests for continuous variables and fisher’s exact tests for categorical variables. The four patients were subselected to restrict the tumor site (three in the right colon, one in the transverse colon), grade (three grade 1, one grade 2), node status (two metastasis cases with N-1a), and account for differences in sex (50% female within both the non-metastasis and metastasis groups). We restricted the cohort to patients without MMR deficiencies as determined through immunohistochemistry (IHC) to control for microsatellite instability status. Tissue blocks were sectioned into 10-micron thick layers, and specific capture areas that contained various distinct macroarchitectural regions (all containing epithelium, tumor-invasive front, intratumoral, lymphatics, etc.) were annotated by the practicing pathologist. A histotechnician carefully extracted / manually cut these capture areas from the tissue, and slides were sent to the Single Cell Genomics Core in the Center for Quantitative Biology for simultaneous H&E staining, imaging, and Visium profiling. After a deparaffinization step, spatial transcriptomics uses spatially-tagged oligo barcodes to 1) register spatial coordinates to collection spots, bound to the mRNA with a poly(A) tail for 2) reverse transcription into cDNA, after 3) permeabilization, and 4) sequencing for mRNA profiling. This allows for unbiased/gridded profiling of up to 5,000 spots (1-10 cells/spot) per 6.5mm by 6.5mm capture area.

### 2.3. Preprocessing

Spatial gene expression profiles contain information for 17,943 genes at almost 5,000 locations per slide (after filtering out non-tissue– total number of Visium dots: 4950, 4922, 4887, and 4169 per slide), sampled in a honeycomb formation. Each Visium spot covers a circular capture area with a diameter of 130 pixels at 20x magnification. After sequencing, we used the *SpaceRanger* package to preprocess the Visium reads into gene count matrices.

As whole slide images (WSIs) derived from the Visium capture areas (size of capture area– 6.5 x 6.5 mm) span thousands of pixels along each dimension, we subdivided the image into square patches centered on the Visium spot. We associated the gene expression of each patch based on the Visium spot at the center of the patch and ignored expression at other spots contained within the patch.

### 2.4. Inference Targets

We used the SpatialDE library to select the top 1000 genes based on their mean fraction of spatial variance (FSV) across all slides (i.e., selected genes which exhibited the greatest spatial variation across the four slides). We tested the capacity of our models to recover expression for all 1000 genes based on dichotomized expression (binary classification) and the original counts (regression) ^16^.

For binary classification, we classify tiles as having a “high” expression for a gene if its individual expression is greater than the median for that slide, as shown by Levy-Jurgensen et al.. For binary tasks, we used a weighted binary cross-entropy loss. The loss was independently taken for positive and negative Visium spots and summed together to account for unbalanced labels.

We model the gene expression distribution for regression tasks as a negative binomial distribution with zero inflation. The model predicts the parameters of the distribution (mean *μ*, dispersion factor *σ*, and inflation of zero count *π*) and is optimized with negative log-likelihood loss. The inferred value is equal to the expected value of the negative binomial distribution (accounting for zero inflation): (1 - *π*)*μ* + *π*(0) = (1 - *π*)*μ*.

### 2.5. Modeling Approaches

First, we compare the performance of the following models using the dichotomized labels and continuous regression objectives. For these models, we sought to establish whether increasing the spatial receptive field by varying the patch size (ablation study) and training graph attention networks (GATs) positively impacted our capacity to predict spatial gene expression. All models featured an output layer, which simultaneously predicted the expression of all selected genes (**Figure 1**). Details of these approaches can be found below:

#### 2.5.1. Local Patch Prediction Model

We initialized our model using InceptionV3 weights (Szegedy et al., 2015) ^17^. InceptionV3 was chosen because it has demonstrated high performance in gene imputation studies by Levy-Jurgensen et al. These models are trained for 25 epochs with a learning rate of 0.0001 and a batch size of 32. The patch size labeled Inception models they were configured to make predictions on (i.e., amount of incorporated surrounding information): **Inception-256** (256 pixels), **Inception-512** (512 pixels), and **Inception-768** (768 pixels) (model suffix indicates patch size).

#### 2.5.2. Contextualized Patch Prediction Models

The Visium spots lie on a hexagonal array, each of which can be treated as a node in a graph, each connected to other Visium spots within 150 pixels. After training the InceptionV3 models from our local patch classification experiments, we extracted patch image embeddings (e.g., n-dimensional descriptive vector) for each Visium spot from the penultimate layer of the trained Inception-256 model. We test a graph attention network (GAT) to see how iterative message-passing can improve model performance. We also compared these results to that obtained using a Vision Transformer (ViT) ^18(p16)^. GAT models were labeled by the number of graph attention layers used to make predictions (i.e., the amount of incorporated surrounding information; the number of layers dictates the size of the neighborhood; eight attention heads per layer): **GAT-1** (1 layer), **GAT-2** (2 layers), and **GAT-4** (4 layers) (model suffix indicates the number of layers). ViT models were labeled by the size of the patch used to form contextual embeddings: **ViT-224** (224-pixel patch sizes) and **ViT-384** (384-pixel patch sizes).

#### 2.5.3. Data Augmentation during training and hyperparameters

To improve the robustness of the model results to new tissue contexts, we transformed patch images by shifting and scaling the pixel intensities by the mean and variance of ImageNet Then, we applied color jitter and random rotations; between −15 and 15 degrees. Random horizontal and vertical flips also augment patches if they are not used in the GAT. Through a coarse hyperparameter search, learning rates were set to 1e-4, and models were trained for 25 epochs. Batch sizes for the Inception model were set to 32, save for the regression models for patch sizes 512 and 768, where batch sizes were set to 16 and 8, respectively.

### 2.6. Comparison of Model Performance

To compare model and patch-size performances, we performed leave-one-slide-out cross-validation (CV), where three of the four slides were used for training/validation, and the final slide was used for testing. This scheme was repeated four times to report on test performance across all four slides in an unbiased manner using macro-averaged (across slides) median (across genes) area under the receiver operating curve (AUROC) and Average Precision (AP) statistics for the binary outcome and correlation coefficients (e.g., Spearman) to compare true versus predicted counts. Non-parametric bootstrapping was used to assess statistical significance through the calculation of a 95% confidence interval. During cross-validation, we use the same set of *InceptionV3* embeddings to train the contextual models (i.e., GAT) that corresponded to the same cross-validation fold. We acknowledge that these statistics could also vary by models, patch size, and slides, so we plotted scatters of test AUCs of each gene to compare these factors in a pairwise manner (e.g., for slide 1, comparing Inception to GAT, or patch size 256 to 768) to draw additional inferences on suitable modeling approaches for a subset of genes. Identifying a subset of genes that obtained optimal performance for one approach versus another was as important as comparing overall performance. These performance differences could suggest relevant tissue features at different scales. For instance, the GAT could extract features from the larger macroarchitecture, indicating its relevance for a gene that does not predict well from *InceptionV3*.

### 2.7. Model Interpretation through Pathway Analysis and Gene Embeddings

Due to fundamental limitations in tissue biology, it is unrealistic to expect that every gene can be predicted from tissue histology. We sought to establish which types of genes could be inferred from histology through pathway analysis. Using the *Elsevier Pathways* database available through the Enrichr package, we performed a pathway analysis of the top 250 genes ranked by AUROC averaged across the CV folds ^19^. Enrichr reports overrepresented pathways using a modified fisher’s exact test. Detected pathways were filtered based on tissue specificity (i.e., could reasonably be involved with the colon). To determine whether different pathways could be inferred from different tissue contexts, pathway analysis results were compared across models.

We also sought to assess how well the predicted gene expression profiles recapitulated relationships/clustering between the Visium spots. This was accomplished through the comparison of Uniform Manifold Approximation and Projection (UMAP) embeddings for the ground truth and predicted expression profiles (on held-out slides) using *InceptionV3* and GAT ^20^. Ground truth and predicted gene expression profiles were projected to a lower dimensional space using AlignedUMAP to preserve the orientation/alignment between the ground truth and predicted expression profiles (i.e., positioning of Visium spots across projections is relatively preserved) to enable comparison between the approaches. Visium spots corresponding to the ground truth UMAP projections were clustered using Hierarchical density-based clustering (HDBSCAN) ^21^. HDBSCAN also identified outlier Visium spots removed from the cluster analysis and scatterplots for all three datasets (ground truth, *InceptionV3*, GAT). Mapper, a Topological Data Analysis method, was used to provide topological summaries of the embeddings by reducing the number of points based on overlap and connectivity ^22–24^. Mapper embedding plots for *InceptionV3* and GAT were colored using the cluster information from the ground truth expression to qualitatively assess topological consistency (i.e., were relationships between Visium spots preserved). This procedure was repeated across the held-out test slides.

### 2.8. Recapitulating Spatial Biology through Spatial Autocorrelation Tests

In addition to tissue clustering, spatial variation is often used as a proxy for explaining the diversity of cellular lineages interacting across the tissue slide. While this factor alone is not an exhaustive assessment of spatial biology, this served as a target for our preliminary assessment (an exhaustive exploration of spatial analyses is out-of-scope of this work, although it is a future direction). After performing cross-validation, we sought to investigate the ability of our algorithms to recapitulate known spatially variable genes on each of the held-out slides from the cross-validation folds. We used two libraries to determine spatially variable (SV) genes: SpatialDE and SPARK-X ^25^. Gene expression counts were summed to create a total count matrix. Genes with low overall expression across the slide (i.e., below a threshold of 30% slide coverage) were removed. The data was then transformed to a normal distribution (by Anscombe transform) to account for the negative binomial distribution of the gene expression. Since SpatialDE’s computation time increases cubically with each additional expression patch, we reduced the resolution of the Visium data through 2×2 median pooling (i.e., taking median expression for specific genes from 2×2 neighborhoods). The reduced memory requirements allowed us to perform further histo-molecular assessments on the images, including clustering based on spatial variability.

Using SpatialDE, we extracted the Fraction of Spatial Variance (FSV) and p-values for each gene from the ground truth set. In addition, we used SpatialDE’s Automatic Expression Histology (AEH) to identify 5 groups of genes that were co-expressed spatially from the ground truth data. Separately, we ran SPARK/SPARK-X on the 1000 genes from each held-out validation slide, reporting projection and adjusted P-value (Bonferroni-corrected) statistics. We separately ran this procedure for the ground truth expression and predicted expression profiles. As an indication of performance, we expected p-values derived for the projection covariance function for both the inferred and original expression data to correlate well with each other for each slide. This was accomplished using the Fisher’s exact test after dichotomizing spatial autocorrelation statistics into “high autocorrelation” (low p-value) and “low autocorrelation” (high p-value) and similar dichotomization for statistics extracted from the inferred expression.

Thresholds for dichotomization were chosen to maximize the magnitudes of the Fisher’s exact test statistics. Dichotomized spatial autocorrelation was cross-tabulated across the genes to report odds ratios with p-values. An odds ratio and corresponding confidence interval of more than one (i.e., statistically significant) would suggest the ability of the model to recapitulate spatial autocorrelation from the slide. Separately, we sought to assess whether genes that were predicted with high accuracy (AUC) were spatially autocorrelated using a similar methodology (i.e., comparing dichotomized spatial autocorrelation on the ground truth expression with dichotomized AUC for each modeling approach). We also compared model accuracies between AEH groups of co-expressed genes identified from SpatialDE using Kruskal-Wallis ANOVA statistical tests. Similar to the AUC comparison, we performed comparisons across models, patch sizes, and slides. It should be noted that these analyses were done on the top 1000 spatially variable genes; dichotomized autocorrelation is for this reference group. For the original and inferred expression, genes which exhibited spatial autocorrelation which differed between patients with and without metastasis were selected for a pathway analysis using the *MSigDB Hallmarks* gene sets via the Enrichr software.

## 3. Results

### 3.1. Model Comparison

First, we assessed the ability of the Inception model to predict gene expression without considering the surrounding tissue macroarchitecture. Across the whole transcriptome, we noted moderately strong concordance between the predicted and actual expression across the held-out slide folds.

#### 3.1.1. Impact of patch size to leverage surrounding spatial context

Importantly, the model achieved optimal performance by increasing access to the surrounding macroarchitecture by increasing the patch size.

#### 3.1.2. Impact of model architecture to leverage surrounding spatial context

After optimizing model architectures, the Inceptionv3 model appeared to outperform the GAT and Transformer approaches for the whole transcriptome assessment. This was true for both classification and regression modeling objectives. The GAT-1 model outperformed the GAT-4 model, which incorporated more of the surrounding tissue context, for the classification task, though GAT-4 outperformed GAT-1 for regression and demonstrated a similar capacity to predict count outcomes as Inception-512 (**Figure 2; Table 1**). A breakdown of these model performances for individual held-out validation slides can be found in the appendix (**Supplementary Table 1; Supplementary Figures 1–2**).

**Figure 2:**
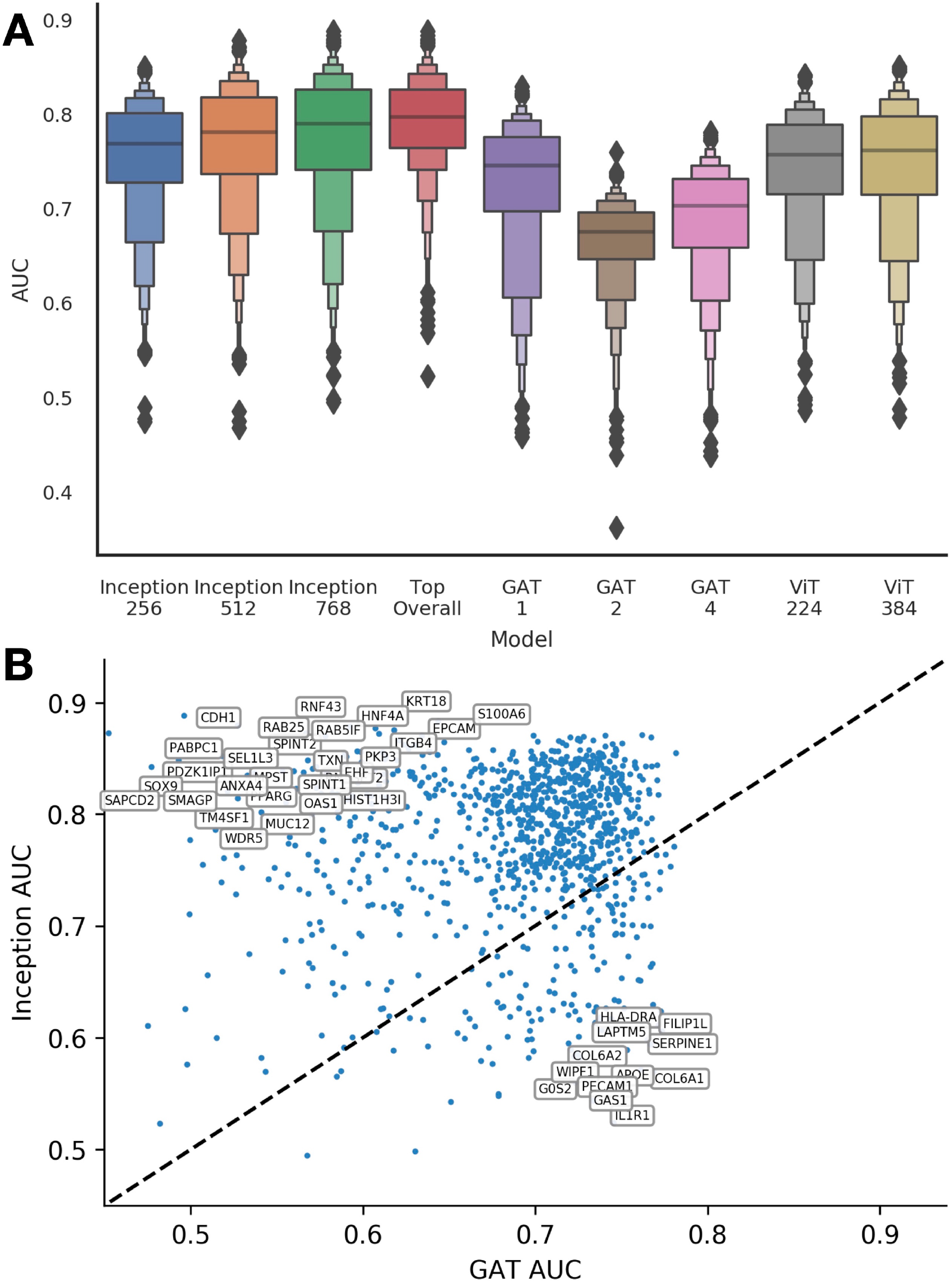
Model Performance Comparison: **A)** Boxenplots which depict the distribution of AUC values for predicting dichotomized expression across all 1000 filtered genes; **B)** Scatter plot of genes representing individual cross-validated AUC values for GAT-4 versus Inception-768; while overall Inception outperforms GAT, there are several genes from which GAT outperformed Inception

**Table 1.** Comparison of model performance for predicting dichotomized and log-transformed expression for genes across the whole transcriptome. These statistics are created by taking the median of each performance statistic across all genes, averaged across held-out slides.

| Model | Classification |  | Regression |  |
| --- | --- | --- | --- | --- |
|  | AUROC | AP | Spearman | Pearson R |
| Inception-256 | 0.769±0.004 | 0.587±0.011 | 0.439±0.011 | 0.466±0.014 |
| Inception-512 | 0.782±0.005 | 0.61±0.013 | 0.456±0.011 | 0.496±0.013 |
| Inception-768 | 0.79±0.006 | 0.621±0.012 | 0.479±0.011 | 0.538±0.013 |
| GAT-1 | 0.746±0.003 | 0.528±0.014 | 0.302±0.009 | 0.309±0.009 |
| GAT-2 | 0.676±0.002 | 0.444±0.01 | 0.409±0.01 | 0.446±0.01 |
| GAT-4 | 0.703±0.004 | 0.459±0.012 | 0.406±0.01 | 0.428±0.011 |
| **ViT-224** | 0.757±0.005 | 0.573±0.012 | 0.432±0.011 | 0.458±0.013 |
| **ViT-384** | 0.762±0.004 | 0.57±0.012 | 0.442±0.012 | 0.473±0.014 |

#### 3.1.3. GAT outperforms Inception on a subset of genes

While overall, Inception outperformed GAT, it is important to recognize that there are many genes for which GAT achieves superior performance (**Figure 2B; Table 2; Supplementary Figure 1**). This was different than comparing multiple Inception models with different patch sizes, where clearly Inception-768 outperformed Inception-256 on nearly all relevant genes (a scatter plot of the specific gene-level AUC for specific genes for patch size 256 compared to 768 can be found in **Supplementary Figure 2**). As different models demonstrate exemplary performance on different subsets of genes, the combined accuracy (AUC) across the genes (i.e., selecting top performing model for each gene based on CV-AUC) is 0.798 (95% CI [0.795-0.802]); the combined AP score is 0.67 (95% CI [0.659-0.677]) (**Figure 2**) (**Top Overall Model**).

**Table 2:**
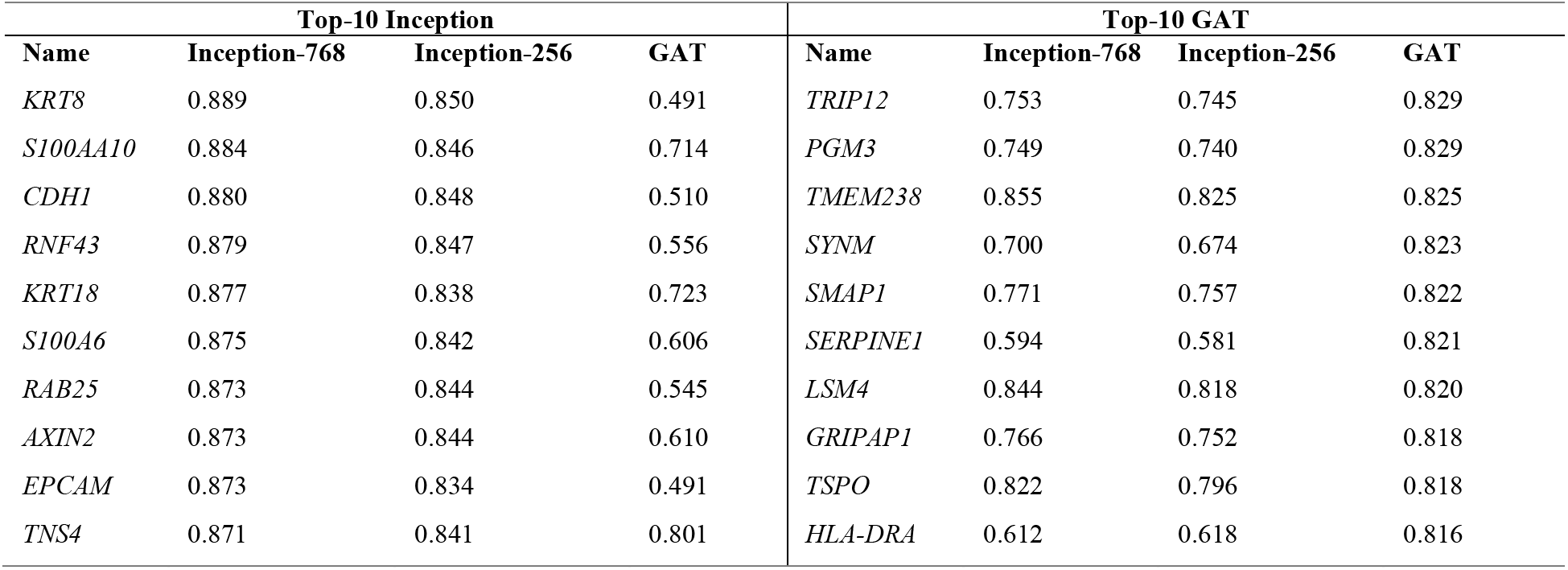
Top 10 performing genes for Inception-768, Inception-256 and GAT-4, ranked by AUC.

### 3.2. Pathway Analysis

The top-performing genes by AUROC for both types of models were also highly related to tumor aggression and migration. For instance, *CDH1*, heavily implicated with tumor suppression, achieved AUROC of 0.880 ^26^. *RAB25* (**Figure 3A**), which can serve as a tumor suppressor or oncogene depending on the context, obtained an AUROC of 0.879. *TNS4* has been heavily implicated for multiple prognostic outcomes following surgery and was predicted with an AUC of 0.871 ^27^. *AXIN2* (**Figure 3C-E**), which inhibits the Wnt signaling pathway and serves to regulate immune cell infiltration ^28^, was detected with an AUROC of 0.873. The pathway analysis corroborated these findings and was highly relevant for metastasis formation. For instance, genes associated with cancer metastasis, cell motility and proliferation, glycolysis, and the epithelium to mesenchymal transition were identified by both models. Interestingly, the GAT model was able to identify genes related to anti-EGFR therapy resistance in colorectal cancer (**Supplementary Table 2**).

**Figure 3.**
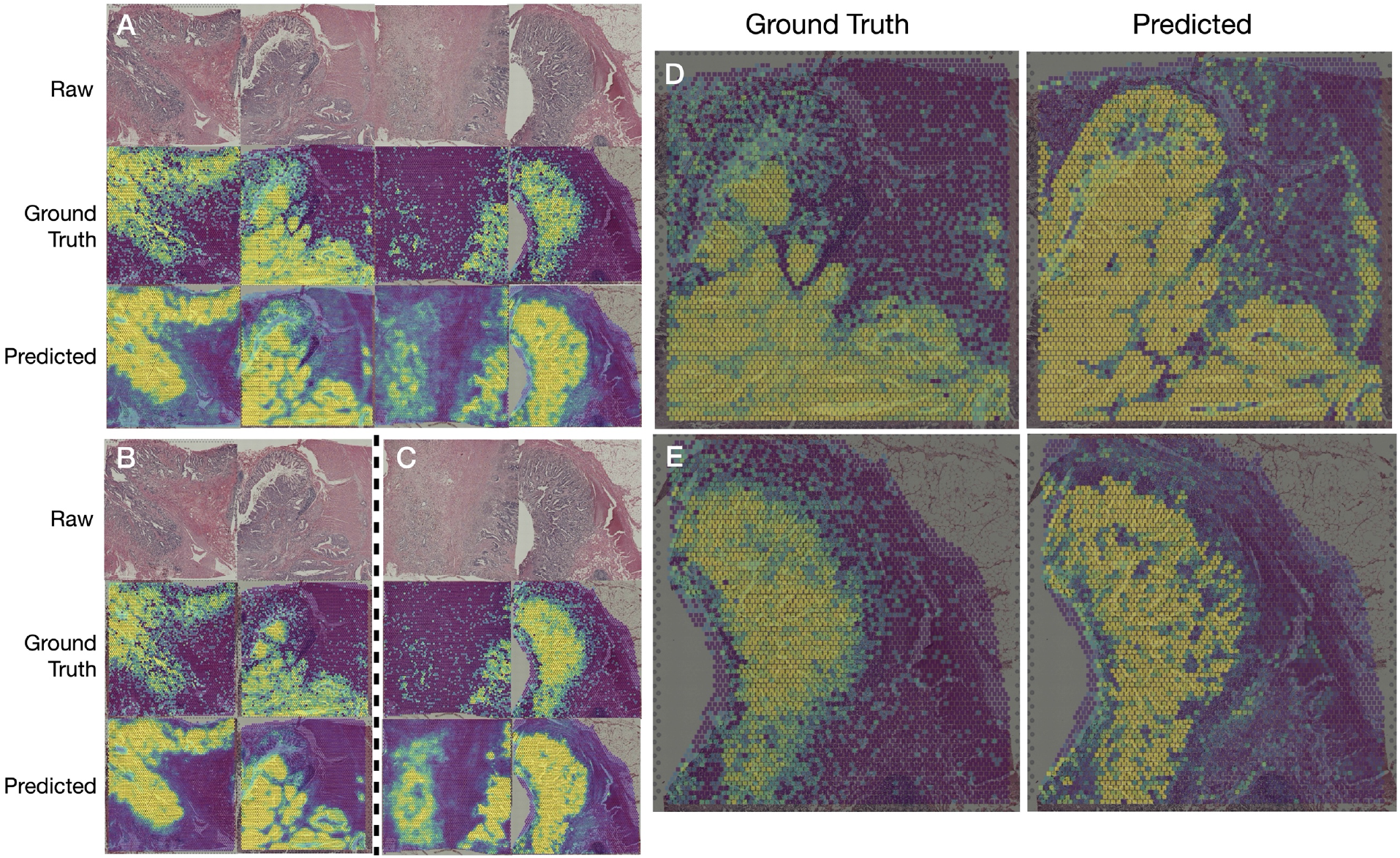
Comparison of log-scaled heatmaps of the ground truth gene expression (top) and output probabilities for above-median expression from the dichotomized classifiers: **A-C)** Inception-768 classifier’s predictions, generating heatmaps for **A)** RAB25, **B)** TNS4, and **C)** AXIN2 genes; all predictions from the held-out test set. The corresponding median AUROCs for these genes were 0.873, 0.871, and 0.873. **D)** AXIN2 imputation on slide B1 with GAT. **E)** AXIN2 imputation on slide D1 with GAT.

#### Inferred Spatial Expression is Topologically Consistent with Ground Truth

Visual inspection of mapper diagrams of clustered UMAP embeddings illustrates topological consistency (i.e., preserve relationships) between the predicted expression and the ground truth (**Figure 4**). Qualitatively, the gene expression embeddings produced from Inception appear to be more closely aligned with the ground truth embedding plots across the tissue types (e.g., **Figure 4 A, C**, where clusters are placed in the approximately same area for Ground Truth and Inception in the Mapper diagrams).

**Figure 4:**
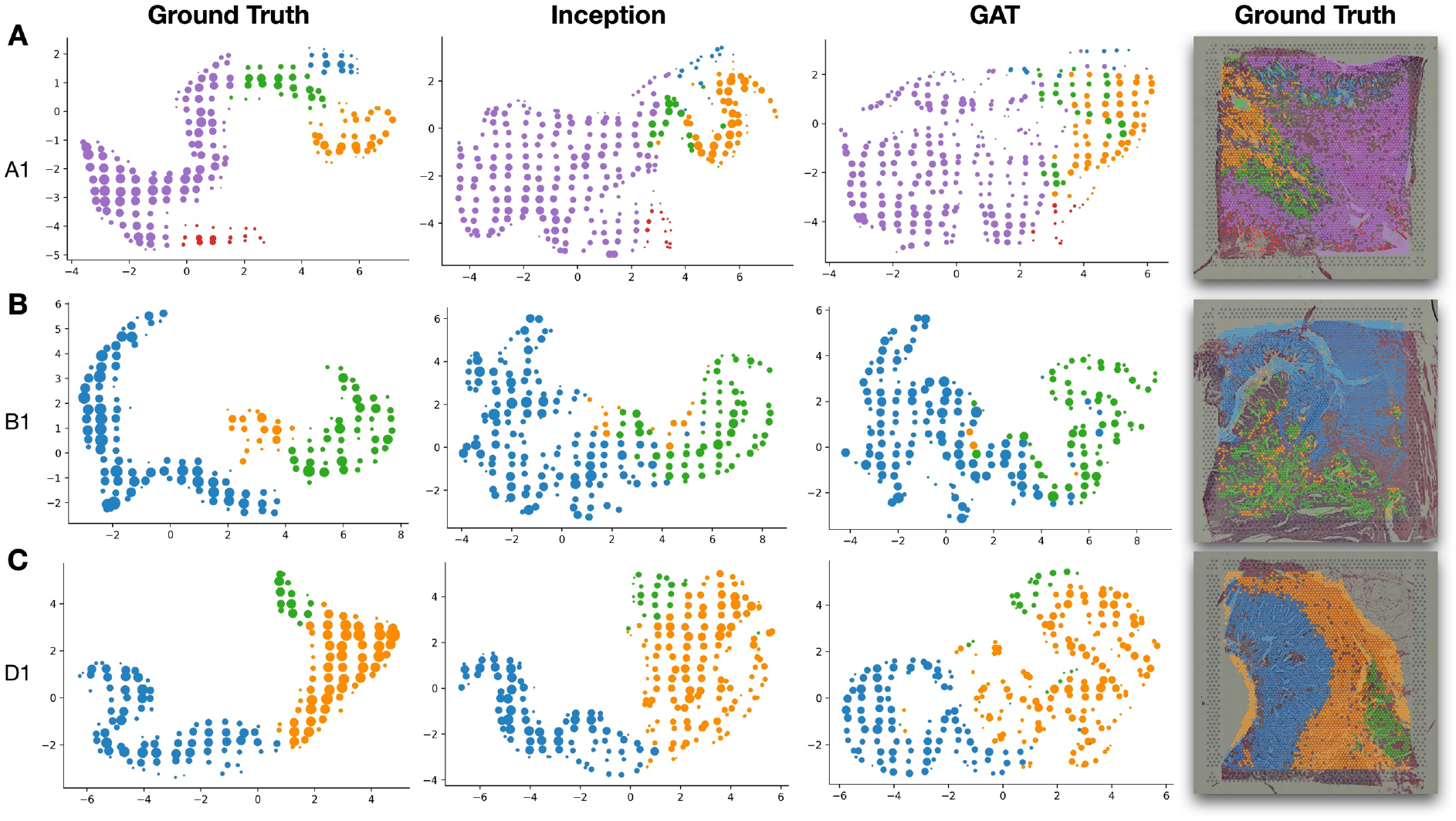
UMAP Embeddings of True and Predicted Gene Expression for slides **A)** A1, **B)** B1 and **C)** D1; embedding plots are summarized using Mapper, which flexibly clusters the expression data with overlapping clusters containing multiple Visium spots; Mapper nodes are sized by the number of associated spots and colored by the dominant cluster in the set of node-associated spots with cluster assignments determined using HDBSCAN; outliers were filtered from these embedding plots and the cluster assignments plotted over the WSI on the right

### 3.3. Spatial Autocorrelation

We also compared the capabilities of each modeling approach for their ability to recapitulate slide-level spatial autocorrelation parameters. Results indicate a large significant positive association between predicted and actual spatial autocorrelation for both Inception and GAT approaches. For half of the held-out slides, GAT demonstrated a larger effect estimate than Inception (**Table 3**).

**Table 3:**
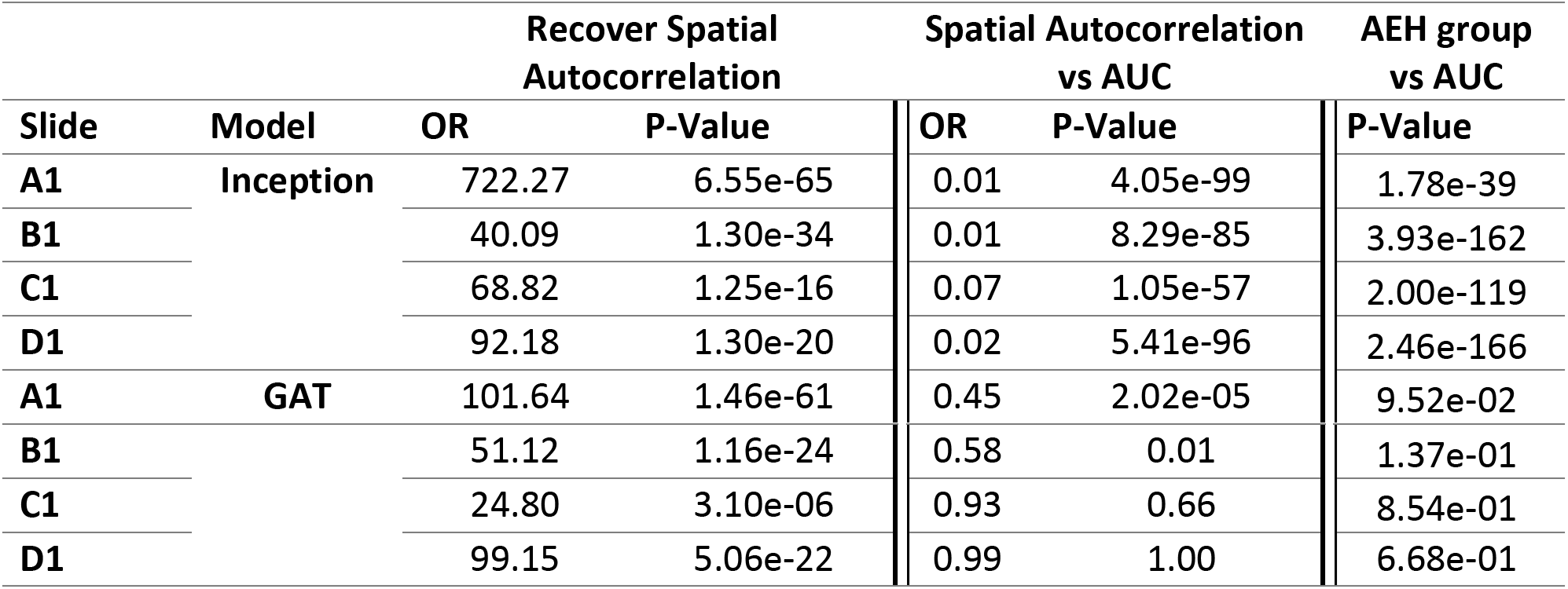
Statistical testing results from spatial autocorrelation analysis, comparing: **A)** Spatial autocorrelation from raw expression (high/low) with inferred spatial autocorrelation from predicted expression values (high/low); **B)** Spatial autocorrelation from raw expression (low/high) with model AUC (high/low); **C)** Whether model accuracy changed depending on the AEH group; **A)** and **B)** were determined through Fisher’s exact tests while **C)** utilized Kruskal-Wallis ANOVA testing

Separately, results indicate that highly spatially autocorrelated genes, as determined using the actual gene expression, were predicted with higher accuracy using Inception and GAT versus genes, which lacked spatial variation (**Table 3**). These models varied in their ability to associate spatial variation with model accuracy. For Inception, there was a large statistically significant effect (**Figure 5**), while spatial variation was not as associated with accuracy for the GAT– i.e., there was a statistically significant association for the first two slides and no statistically significant effect for the final two. For the Inception model, groups of genes that tended to be co-expressed, as determined through the AEH analysis on the raw expression data, were found to have widely different accuracies (**Table 3; Supplementary Figures 3–5**). Similar to the spatial variation analysis, GAT model accuracy did not vary substantially between AEH groups determined from the raw expression data.

**Figure 5.**
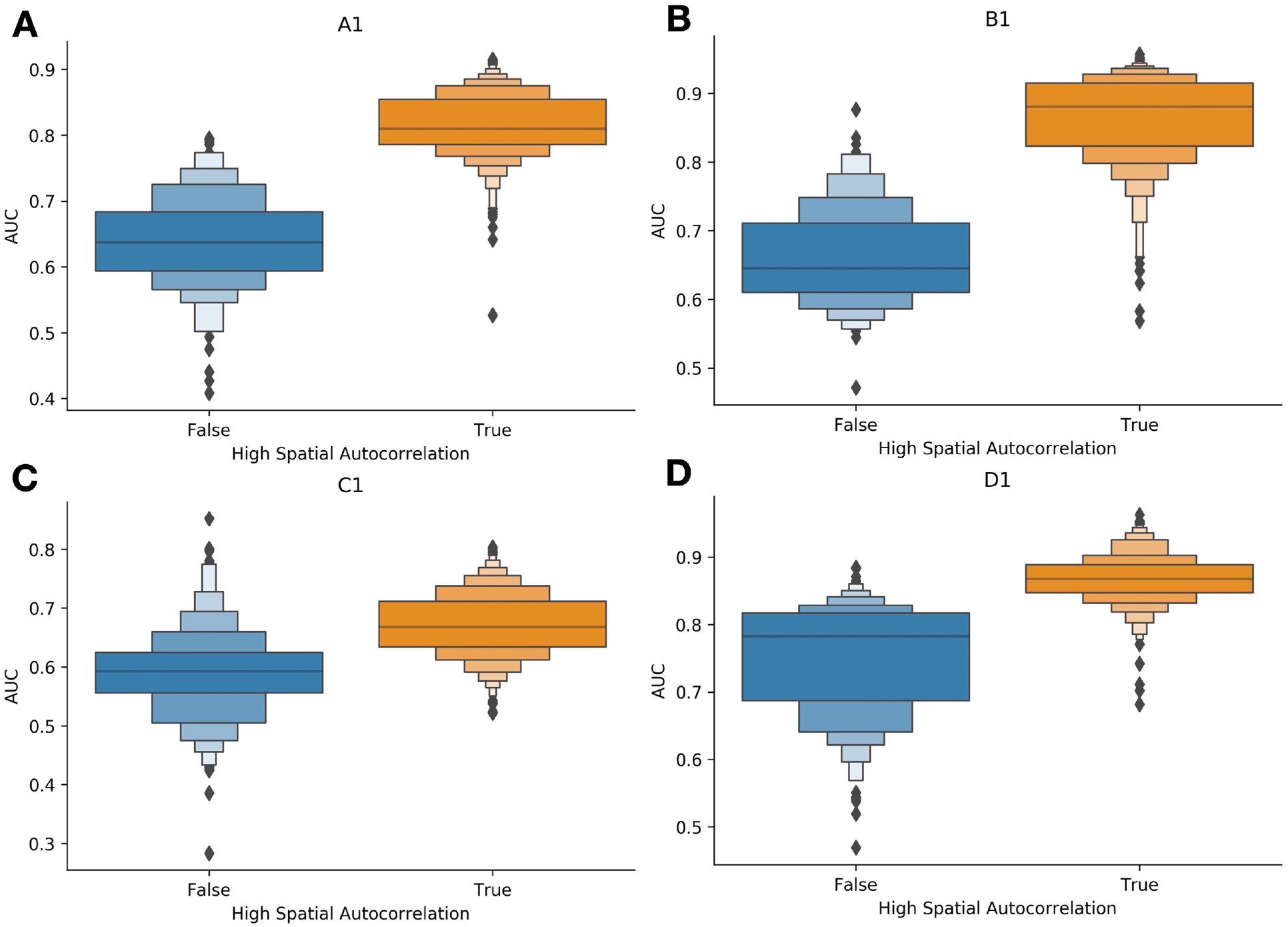
Boxenplots illustrating the predictive accuracy of Inceptionv3 (AUC, y-axis) across genes, separated by whether highly significant spatial variation was reported (blue versus orange, x-axis); gathered from validation slides held-out of the training/validation set. The positive association demonstrates higher accuracy for genes with significant spatial variability (i.e., not distributed randomly); these genes were more accurately predicted by our RNA inference model. **A-D)** correspond to slides A1-D1, respectively

Using the ground truth, Inception, and GAT results, we identified the set of genes, which exhibited different spatial variation for primary sites with (A1, B1) and without metastasis (C1, D1) based on the dichotomized thresholds. Through a pathway analysis (gene set testing of Cancer Hallmark genes), genes that were differently autocorrelated were related to the epithelial to mesenchymal transition (**Supplementary Table 3**).

## 4. Discussion

Assessment of Colon cancer tumor recurrence risk depends on the ability to assess lymph node status. For cases where lymph nodes are unable to be completely assessed, leveraging information found in the tumor immune microenvironment at the primary resection site presents a viable alternative. Yet most technologies to assess the primary site lack a spatial component (e.g., bulk expression), which does not enable a comprehensive characterization of the tissue. Spatial transcriptomic technologies enable high multiplexing at incredible spatial resolution. Due to both fiscal and sampling (i.e., placement of capture area) constraints, findings are not likely to be clinically actionable or reproducible through a low-cost test. Inferring a spatial digital biomarker from a routine histological slide (WSI) has the potential to enable high-throughput tissue characterization for the creation of nascent decision-making aids which can complement existing specimen findings. In this study, we explored the potential for deep learning models to predict spatial gene expression from formalin-fixed specimens, the ability to recapitulate well known spatial findings, and comment on the degree to which spatial information at higher-order contexts (i.e., macroarchitecture) plays a role in modeling spatial expression.

We add further evidence that deep learning models can accurately predict gene expression in whole slide images. Furthermore, we demonstrate that increasing the receptive field can improve the performance of certain subsets of genes. Interestingly, although InceptionV3 outperformed other modeling approaches overall, this did not apply to all genes. We noted that certain genes were predicted more effectively at local spatial resolutions (Inception) while others benefited from considering a broader architectural context (GAT). For instance, we noticed that *COL6A1* and *COL6A2* were consistently predicted better by the GAT model as compared to Inceptionv3 ^29–31^. *COL6A1* is a crucial component of collagen secretion, extracellular matrix maintenance, and mesenchymal phenotype promotion. While both models demonstrated the capacity to recapitulate well-known cancer markers, it was clear that certain genes can be better predicted by considering receptive field size and choosing a model that best incorporates this spatial information.

Spatial autocorrelation was recapitulated for these slides from the deep learning model predictions. We noticed that increases in modeling accuracy were associated with spatial gene expression variation. Genes that were differentially autocorrelated between metastatic and non-metastatic tumors corroborated with well-established oncogenic pathways. This supports our overarching modeling approach to characterize spatial heterogeneity. As some genes were better predicted using the broader spatial architecture, it would make sense to decide which modeling approach to utilize on a gene-by-gene basis and whether the local versus spatial context is preferred. This allows for the selection of optimal modeling approaches in a gene-specific manner to extend the broad applicability of our framework across all disease-relevant genes.

Importantly, the techniques featured in this work will prove useful for inferring gene expression on slides from external cohorts. This is necessary due to the prohibitive costs of spatial transcriptomics. If validated properly on a slightly larger set of slides while carefully controlling for potential confounders (e.g., site, MMR status, etc.), these methods may overcome sources of patient-specific batch effects to allow the report of less biased effect estimates for large-scale cohort studies. Validation efforts on these held-out slides can feature validation assessments such as Spark-X and SpatialDE to ensure spatial heterogeneity from the inferred expression data is similar to the initial internal validation cohort. To this end, we achieved remarkable performance for the ability to predict spatial heterogeneity, which will potentially help power future studies elucidating spatial transcriptomic predictors of colon metastasis.

Our results did not support the hypotheses that, overall, spatial gene expression estimation would benefit from message-passing between patch locations via transformer models and graph convolutional networks. However, these neural networks outperform local patch prediction across a large set of genes, suggesting the relevance of these methods for genes which leverage the spatial context. For GAT, underperforming genes can potentially be explained by the fact that graph convolutional networks tend to smoothen features (Li et al., 2018), which may weaken the model’s predictions if the gene expression data is not as smooth. While these findings could alternatively suggest that neighboring patches may not be as correlated as suspected, this may only hold true for a subset of genes. Increasing graph convolution layers may result in optimal performance for genes, which rely on higher-order dependencies between tissue regions; however, such genes may be outnumbered by genes better suited for the Inception approach.

There were a few study limitations of note. Our validation scheme featured the use of held-out slides. Generally, as two cases had tumor metastasis, whereas the other two did not have concurrent metastasis, we believed held-out slides would have similar heterogeneity in expression and morphology as the training/validation slides. Indeed, it is possible that tissue expression/morphology existed outside of this range, differentially impacting the GAT/transformer models designed to capture long-range spatial dependencies. As we had manually selected capture areas, we did not expect slides to have exactly matching/analogous architectural features; thus, the GAT/transformer model could have been unable to generalize as it aims to integrate this higher-level information. There were four whole slide images for both training and cross-validation, and while these provided over ten thousand patches in the training set when taken individually, they provided limited opportunities for models to learn diverse global contexts that transformer models would have benefitted from. These challenges could have been ameliorated by pretraining the GAT/Transformer on a variety of colorectal cancer tissue contexts, which does not explicitly require spatial omics data. We also acknowledge the impact of evolving workflows for spatial resolution of transcriptomics information. Many workflows rely on the prediction of tissue features from fresh or fresh frozen tissue. We utilized formalin-fixed paraffin-embedded (FFPE) tissue slides. Only recently have deparaffinization and assay workflows been developed for FFPE. As many of these specimen processing workflows are still under development, manual staining and imaging could have both introduced batch effects and impacted tissue quality. For instance, while a plethora of data preprocessing workflows offer capabilities to combat batch effects through normalization, a robust comparison of preprocessing workflows was outside of the study scope. Our group is keen to adopt future iterations of the FFPE workflows. We expect that protocol updates for the FFPE workflow and data collection will provide major improvements in specimen processing, data quality, and resolution, improving prediction models. Enlarging the capture area and evaluating complementary molecular assays (e.g., spatial proteomics) will improve the resolution and scope of our findings. Although this work does not explicitly assess the tumor immune microenvironment, the methodology explored here can help facilitate spatial analyses and will be used to motivate future clinical findings. Future work will clearly demarcate these regions (e.g., intratumoral, invasive margin, away from the tumor) for a more granular analysis that is well-adjusted for potential confounders.

## 5. Conclusions

Tumor metastasis is heavily tied to poor prognostic outcomes and risk of recurrence. Assessment of key niches at the primary site (e.g., tumor immune microenvironment) may reveal histomorphological or biological factors relating to nodal involvement or distant metastasis. In this work, we investigated the potential to infer spatially resolved transcriptomics from the tissue histology of colorectal cancer patients through several sophisticated neural network approaches. Our findings indicate that neural networks can be effectively employed in this capacity and that selection of a neural network model could be informed by its relevance to a molecular pathway resolved from histological features at different scales. Findings reaffirm the role of the epithelial-to-mesenchymal transition as an important metastasis-related process. In the future, with additional algorithmic fine-tuning, data curation, and standardization, there are opportunities to generalize these findings to perform large-scale spatial molecular assessments.

## Authors’ Contributions

The conception and design of the study were contributed by JL and LV. MF, EF, and CS spearheaded study design, analysis, interpretation and initial manuscript drafting. All authors contributed to writing and editing of the manuscript and all authors read and approved the final manuscript.

## Acknowledgements

We would like to thank Gabriel Brooks for his thoughtful discussion of the subject matter. This work was supported by NIH grants R01CA216265, R01CA253976, and P20GM104416 to BC, Dartmouth College Neukom Institute for Computational Science CompX awards to BC, JL and LV, and DCC, DPLM Clinical Genomics and Advanced Technologies EDIT program. JL is supported through NIH subawards P20GM104416 and P20GM130454. The funding bodies above did not have any role in the study design, data collection, analysis and interpretation, or writing of the manuscript.

## Appendix

**Supplementary Table 1.** Comparison of model performance for predicting dichotomized and log-transformed expression for genes across the whole transcriptome. These statistics are created by taking the median across all genes, reported for each held-out slide.

| Model | Slide | Classification |  | Regression |  |
| --- | --- | --- | --- | --- | --- |
|  |  | AUROC | AP | Spearman R | Pearson R |
| Inception-256 | Overall | 0.768 | 0.603 | 0.444 | 0.477 |
|  | A1 | 0.78 | 0.675 | 0.494 | 0.511 |
|  | B1 | 0.834 | 0.796 | 0.612 | 0.605 |
|  | C1 | 0.625 | 0.432 | 0.214 | 0.301 |
|  | D1 | 0.831 | 0.51 | 0.454 | 0.491 |
| Inception-512 | Overall | 0.781 | 0.63 | 0.461 | 0.498 |
|  | A1 | 0.79 | 0.688 | 0.515 | 0.537 |
|  | B1 | 0.856 | 0.816 | 0.639 | 0.647 |
|  | C1 | 0.629 | 0.459 | 0.22 | 0.299 |
|  | D1 | 0.847 | 0.555 | 0.469 | 0.51 |
| Inception-768 | Overall | 0.787 | 0.639 | 0.486 | 0.54 |
|  | A1 | 0.802 | 0.705 | 0.551 | 0.578 |
|  | B1 | 0.864 | 0.823 | 0.671 | 0.655 |
|  | C1 | 0.636 | 0.474 | 0.239 | 0.4 |
|  | D1 | 0.845 | 0.553 | 0.481 | 0.525 |
| GAT-1 | Overall | 0.743 | 0.539 | 0.413 | 0.43 |
|  | A1 | 0.785 | 0.669 | 0.364 | 0.376 |
|  | B1 | 0.811 | 0.73 | 0.603 | 0.582 |
|  | C1 | 0.606 | 0.409 | 0.239 | 0.335 |
|  | D1 | 0.77 | 0.349 | 0.444 | 0.428 |
| GAT-2 | Overall | 0.677 | 0.444 | 0.407 | 0.45 |
|  | A1 | 0.769 | 0.645 | 0.511 | 0.545 |
|  | B1 | 0.772 | 0.612 | 0.574 | 0.568 |
|  | C1 | 0.589 | 0.333 | 0.146 | 0.265 |
|  | D1 | 0.578 | 0.184 | 0.421 | 0.428 |
| GAT-4 | Overall | 0.702 | 0.462 | 0.453 | 0.499 |
|  | A1 | 0.679 | 0.571 | 0.517 | 0.551 |
|  | B1 | 0.764 | 0.614 | 0.622 | 0.643 |
|  | C1 | 0.604 | 0.379 | 0.228 | 0.356 |
|  | D1 | 0.759 | 0.283 | 0.445 | 0.454 |
| GAT-4-AXIN2 | Overall | 0.702 | 0.462 | 0.545 | 0.578 |
|  | A1 | 0.679 | 0.57 | 0.511 | 0.446 |
|  | B1 | 0.764 | 0.614 | 0.745 | 0.771 |
|  | C1 | 0.604 | 0.379 | 0.286 | 0.46 |
|  | D1 | 0.759 | 0.283 | 0.637 | 0.633 |
| GAT-4-CDH1 | Overall | 0.800 | 0.707 | 0.546 | 0.571 |
|  | A1 | 0.826 | 0.747 | 0.468 | 0.478 |
|  | B1 | 0.858 | 0.791 | 0.753 | 0.741 |
|  | C1 | 0.648 | 0.462 | 0.31 | 0.425 |

|  | D1 | 0.867 | 0.827 | 0.652 | 0.64 |
| --- | --- | --- | --- | --- | --- |
| **GAT-1-ZINB** | Overall | n/a | n/a | 0.393 | 0.408 |
|  | A1 | n/a | n/a | 0.441 | 0.429 |
|  | B1 | n/a | n/a | 0.551 | 0.562 |
|  | C1 | n/a | n/a | 0.15 | 0.19 |
|  | D1 | n/a | n/a | 0.429 | 0.449 |
| **GAT-4-ZINB** | Overall | n/a | n/a | 0.41 | 0.432 |
|  | A1 | n/a | n/a | 0.485 | 0.542 |
|  | B1 | n/a | n/a | 0.602 | 0.621 |
|  | C1 | n/a | n/a | 0.128 | 0.2 |
|  | D1 | n/a | n/a | 0.426 | 0.364 |
| **ViT-224** | Overall | 0.756 | 0.583 | 0.436 | 0.454 |
|  | A1 | 0.768 | 0.644 | 0.511 | 0.506 |
|  | B1 | 0.826 | 0.762 | 0.601 | 0.567 |
|  | C1 | 0.604 | 0.413 | 0.177 | 0.255 |
|  | D1 | 0.827 | 0.513 | 0.455 | 0.487 |
| **ViT-384** | Overall | 0.765 | 0.601 | 0.446 | 0.468 |
|  | A1 | 0.794 | 0.678 | 0.513 | 0.521 |
|  | B1 | 0.836 | 0.788 | 0.612 | 0.583 |
|  | C1 | 0.606 | 0.406 | 0.213 | 0.307 |
|  | D1 | 0.823 | 0.530 | 0.446 | 0.462 |

**Supplementary Figure 1:**
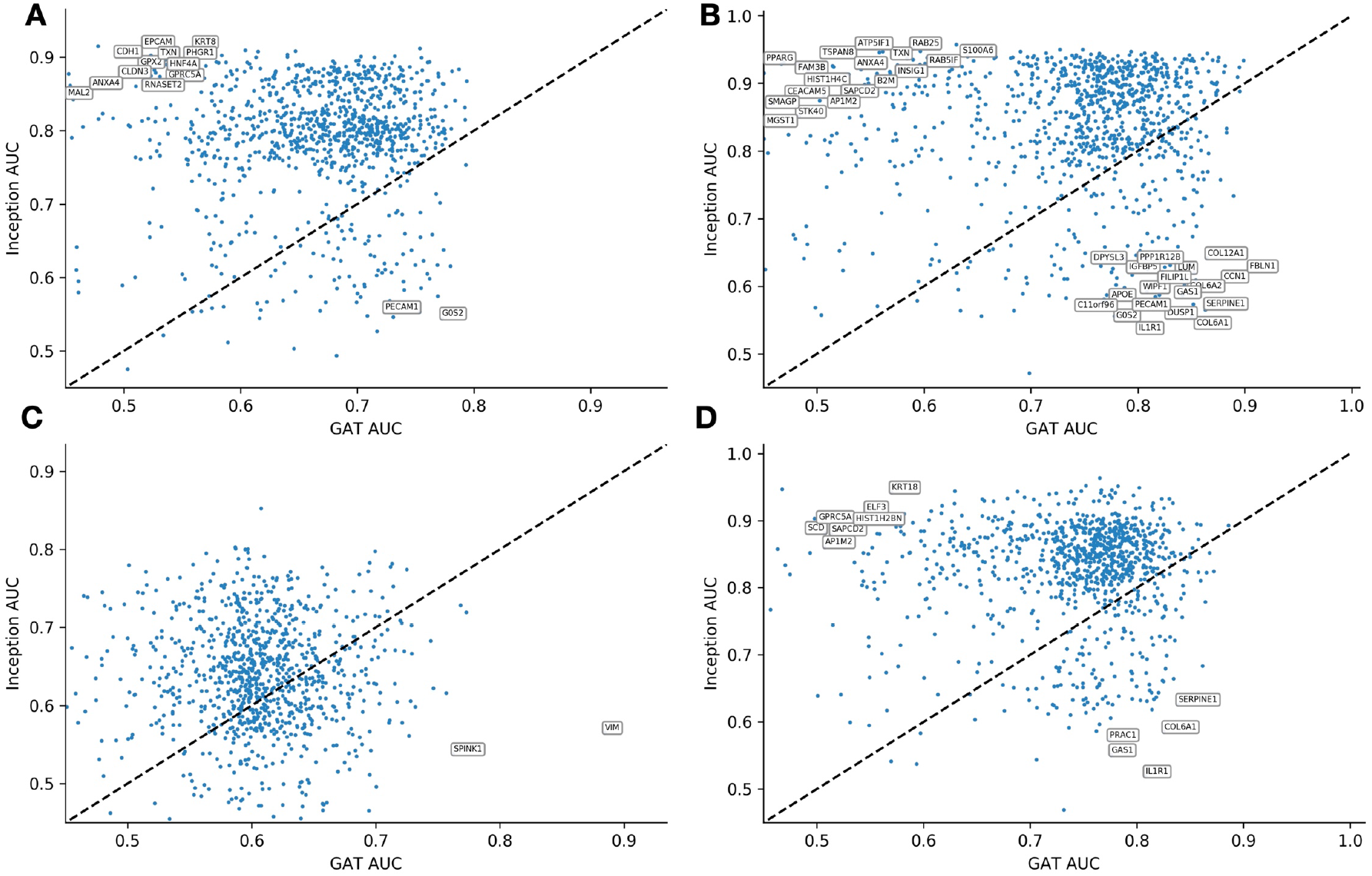
Scatterplot depicting gene-specific AUCs for GAT and Inception-768 for slides: **A)** A1; **B)** B1; **C)** C1; **D)** D1

**Supplementary Figure 2:**
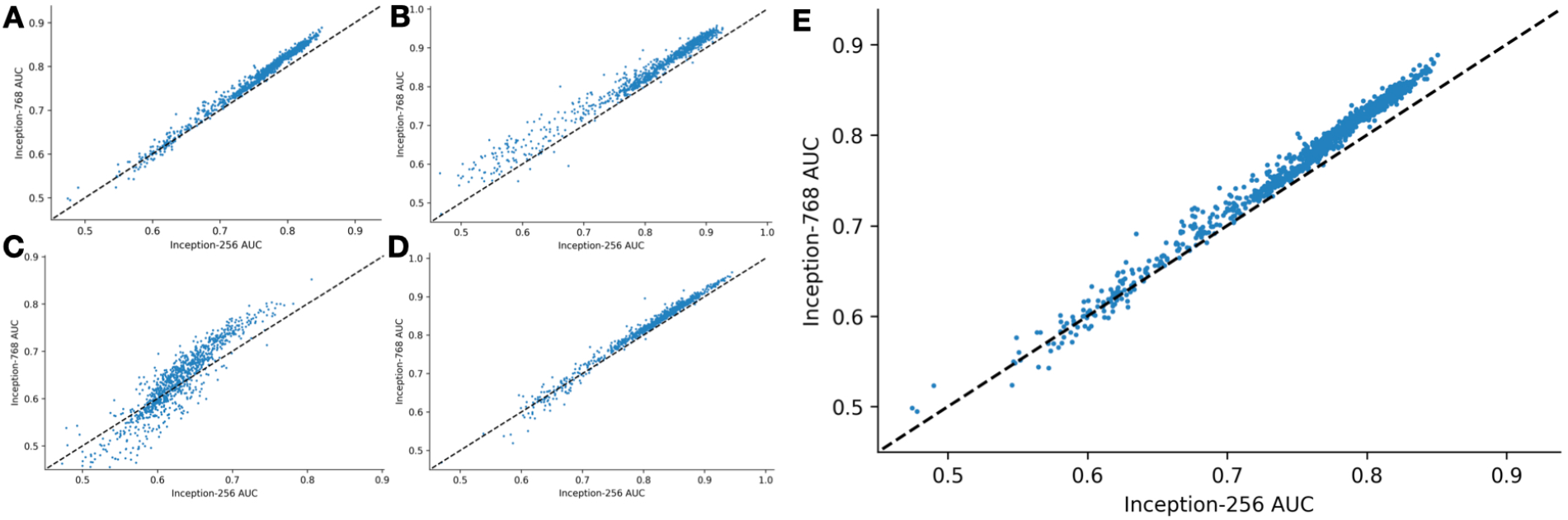
Scatterplot depicting gene-specific AUCs for Inception-256 and Inception-768 for slides: **A)** A1; **B)** B1;**C)** C1; **D)** D1, **E)** Overall

**Supplementary Table 2:**
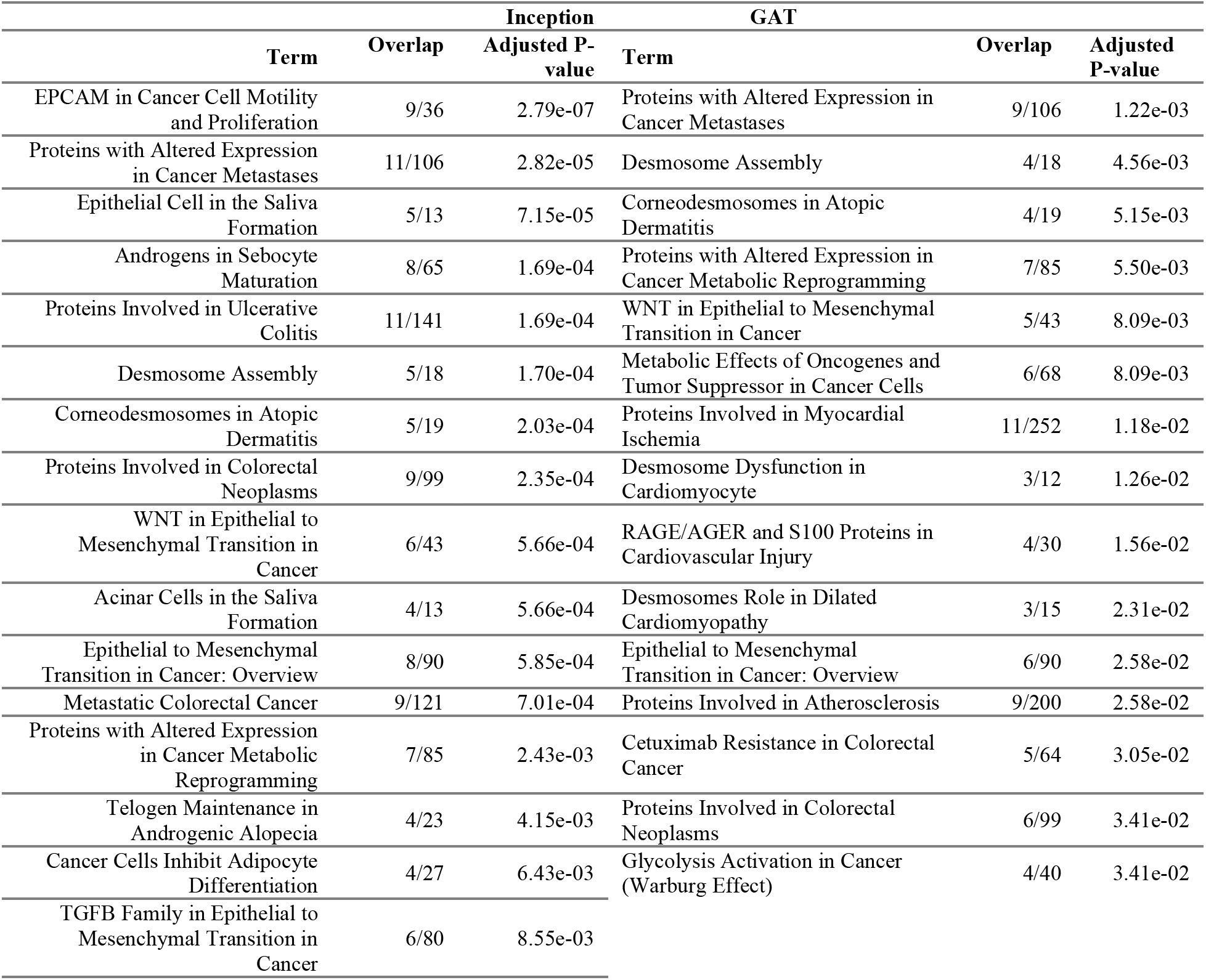
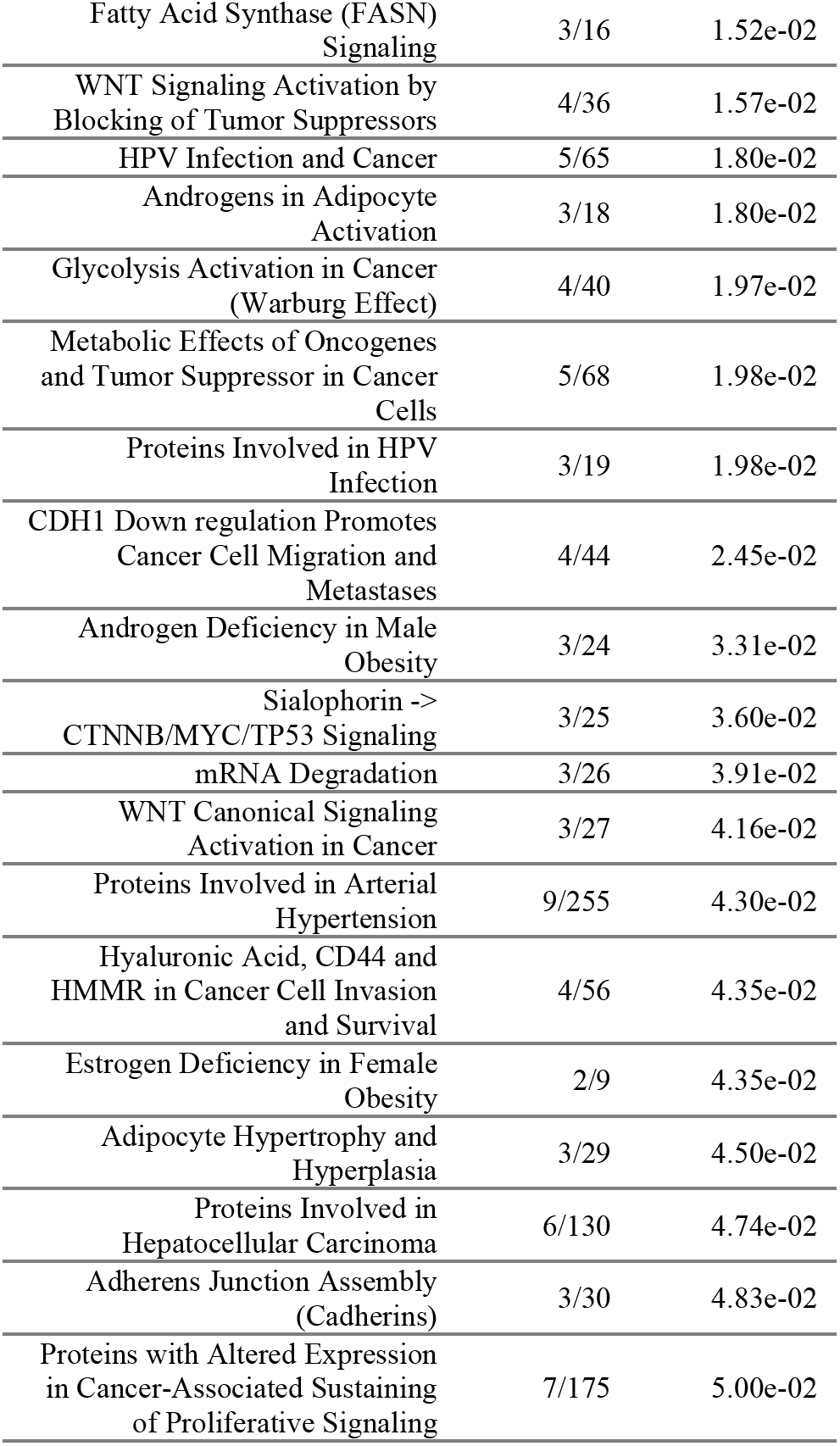
Enrichr pathway results for genes found to be accurately predicted from the tissue histology via the Inception and GAT approaches; pathways were filtered based on tissue specificity

**Supplementary Figure 3:**
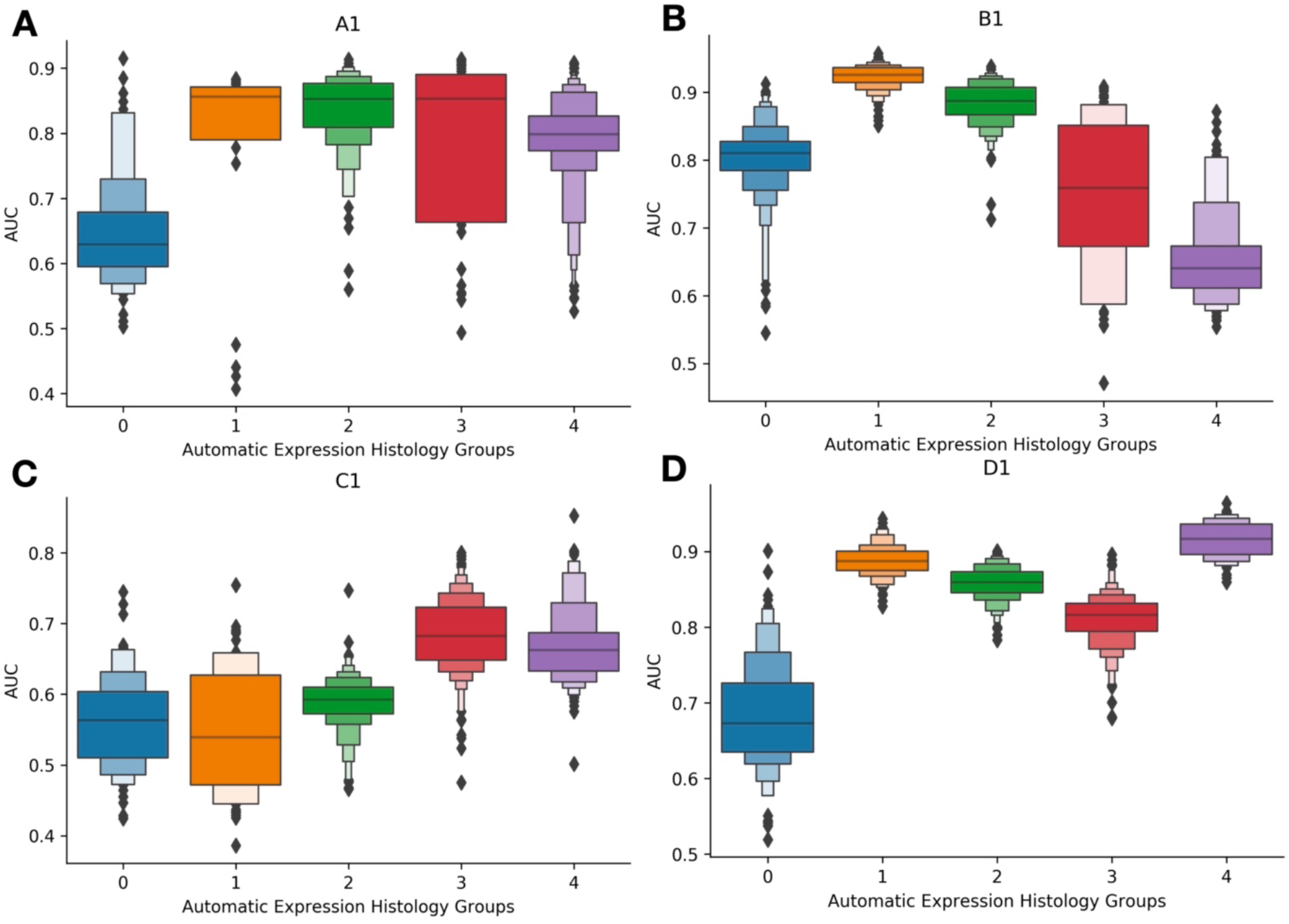
Boxenplots illustrating the predictive accuracy of Inceptionv3 (AUC, y-axis) across genes, separated by the genes’ Automatic Expression Histology groups (colors, x-axis); gathered from validation slides held-out of the training/validation set. Different groups were assigned for each slide. **A-D)** correspond to slides A1-D1 respectively

**Supplementary Figure 4:**
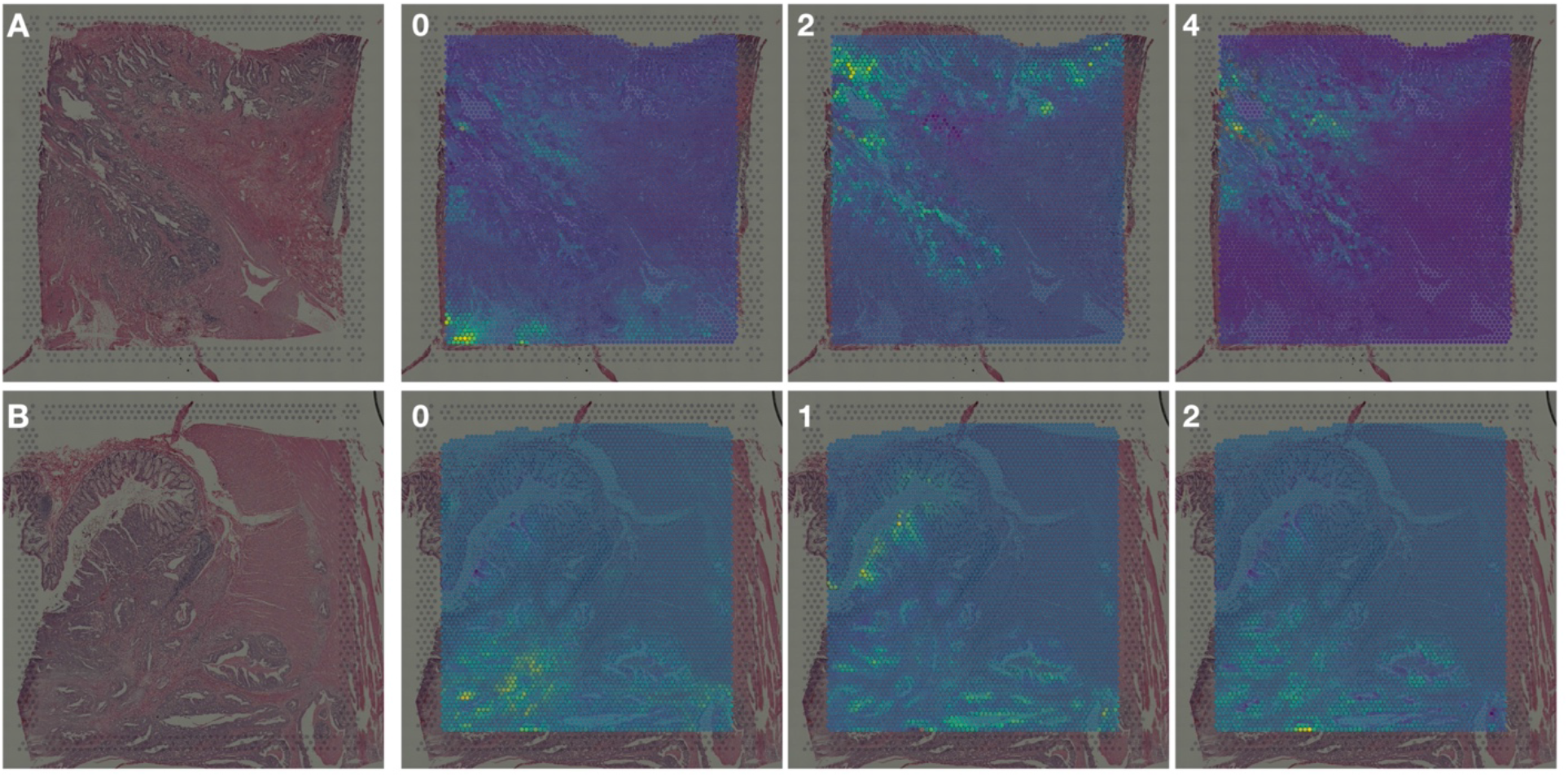
Spatial expression patterns for select AEH groups for slides: **A)** A1; **B)** B1; genes from the first featured AEH group for each slide were predicted with low accuracy; genes from the final two AEH groups were predicted with high accuracy

**Supplementary Figure 5:**
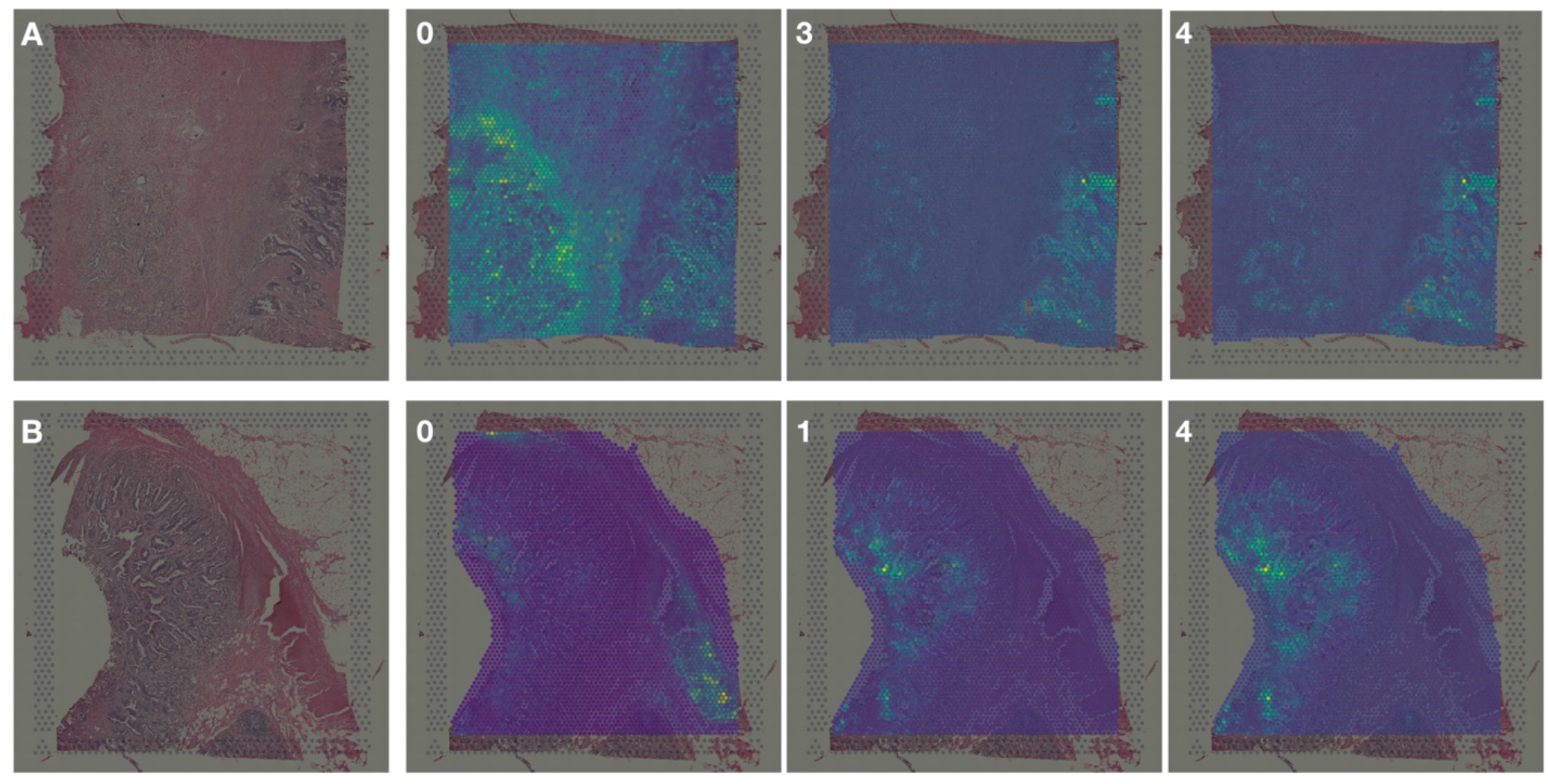
Spatial expression patterns for select AEH groups for slides: **A)** C1; **B)** D1; genes from the first featured AEH group for each slide were predicted with low accuracy; genes from the final two AEH groups were predicted with high accuracy

**Supplementary Table 3:** Genes differentially spatially autocorrelated between METS/No-METS.

| Ground Truth |  |  |  |
| --- | --- | --- | --- |
| Randomly Selected Genes | Pathways | Overlap | Adj. P-Value |
| STXBP3 | Epithelial Mesenchymal Transition | 17/200 | 2.53e-09 |
| IMPAD1 | Coagulation | 11/138 | 0.000006432 |
| ING1 | TGF-beta Signaling | 11/200 | 0.002024 |
| YY1AP1 | Myogenesis | 11/200 | 0.0001214 |
| IGFBP4 | Apical Junction | 10/200 | 0.0001214 |
| ACO1 | Hypoxia | 5/54 | 0.0005478 |
| ID3 | Cholesterol Homeostasis | 6/144 | 0.02979 |
| PTAR1 | UV Response Dn | 7/199 | 0.02582 |
| SYNM | UV Response Up | 7/200 | 0.0286 |
| SPRY2 | IL-2/STAT5 Signaling | 7/200 | 0.02582 |
| RABL3 | Estrogen Response Late | 6/158 | 0.02582 |
| N4BP2L2 | p53 Pathway | 4/74 | 0.02582 |
| Inception |  |  |  |

| Randomly Selected Genes | Pathways | Overlap | Adj. P-Value |
| --- | --- | --- | --- |
| DPYSL3 | Epithelial Mesenchymal Transition | 9/200 | 3.41e-13 |
| CALD1 | TGF-beta Signaling | 3/138 | 0.004267 |
| G0S2 | Coagulation | 3/200 | 0.001833 |
| IGHG1 | TNF-alpha Signaling via NF-kB | 2/54 | 0.003624 |
| LIMS2 | Angiogenesis | 2/200 | 0.05432 |
| COL6A2 | Xenobiotic Metabolism | 2/200 | 0.02452 |
| C1S | Hypoxia | 2/200 | 0.02452 |
| SERPINF1 | Myogenesis | 2/200 | 0.02452 |
| FOSB | Complement | 2/200 | 0.02452 |
| SORBS1 | Inflammatory Response | 1/36 | 0.02452 |
| WIPF1 | IL-6/JAK/STAT3 Signaling | 1/87 | 0.1169 |
| PCOLCE | Interferon Alpha Response | 1/97 | 0.119 |

GAT
| Randomly Selected Genes | Pathways | Overlap | Adj. P-Value |
| --- | --- | --- | --- |
| ACTA2 | Epithelial Mesenchymal Transition | 18/200 | 1.85e-27 |
| COL5A2 | Angiogenesis | 4/36 | 0.000002955 |
| COL3A1 | Myogenesis | 5/200 | 0.00009592 |
| COL12A1 | Glycolysis | 3/200 | 0.01585 |
| GREM1 | Apical Junction | 3/200 | 0.01585 |
| CDH11 | Hedgehog Signaling | 2/138 | 0.1196 |
| SPARCL1 | Apical Surface | 2/144 | 0.1321 |
| CRYAB | Coagulation | 2/200 | 0.06564 |
| POSTN | UV Response Dn | 2/200 | 0.06564 |
| TPM2 | TGF-beta Signaling | 1/36 | 0.1475 |
| DES | Hypoxia | 1/44 | 0.09354 |
| DCN | Adipogenesis | 1/54 | 0.09354 |

## References

1. Wong MC, Huang J, Lok V, et al. Differences in incidence and mortality trends of colorectal cancer worldwide based on sex, age, and anatomic location. Clin Gastroenterol Hepatol. 2021;19(5):955–966.

2. Amin MB, Edge S, Greene F, et al., eds. AJCC Cancer Staging Manual. 8th ed. Springer International Publishing; 2017. Accessed April 28, 2021. https://www.springer.com/gp/book/9783319406176

3. Senthil M, Trisal V, Paz IB, Lai LL. Prediction of the Adequacy of Lymph Node Retrieval in Colon Cancer by Hospital Type. Arch Surg. 2010;145(9):840–843. doi:10.1001/archsurg.2010.182

4. Nearchou IP, Gwyther BM, Georgiakakis ECT, et al. Spatial immune profiling of the colorectal tumor microenvironment predicts good outcome in stage II patients. Npj Digit Med. 2020;3(1):1–10. doi:10.1038/s41746-020-0275-x

5. Uttam S, Stern AM, Sevinsky CJ, et al. Spatial domain analysis predicts risk of colorectal cancer recurrence and infers associated tumor microenvironment networks. Nat Commun. 2020;11(1):3515. doi:10.1038/s41467-020-17083-x

6. Binnewies M, Roberts EW, Kersten K, et al. Understanding the tumor immune microenvironment (TIME) for effective therapy. Nat Med. 2018;24(5):541–550. doi:10.1038/s41591-018-0014-x

7. Bruni D, Angell HK, Galon J. The immune contexture and Immunoscore in cancer prognosis and therapeutic efficacy. Nat Rev Cancer. 2020;20(11):662–680. doi:10.1038/s41568-020-0285-7

8. Wu Y, Cheng Y, Wang X, Fan J, Gao Q. Spatial omics: Navigating to the golden era of cancer research. Clin Transl Med. 2022;12(1):e696. doi:10.1002/ctm2.696

9. Suwalska A, Zientek L, Polanska J, Marczyk M. Quantifying Spatial Heterogeneity of Tumor-Infiltrating Lymphocytes to Predict Survival of Individual Cancer Patients. J Pers Med. 2022;12(7):1113. doi:10.3390/jpm12071113

10. LeCun Y, Bengio Y, Hinton G. Deep learning. Nature. 2015;521(7553):436–444. doi:10.1038/nature14539

11. He B, Bergenstråhle L, Stenbeck L, et al. Integrating spatial gene expression and breast tumour morphology via deep learning. Nat Biomed Eng. Published online June 22, 2020:1–8. doi:10.1038/s41551-020-0578-x

12. Levy-Jurgenson A, Tekpli X, Kristensen VN, Yakhini Z. Spatial transcriptomics inferred from pathology whole-slide images links tumor heterogeneity to survival in breast and lung cancer. Sci Rep. 2020;10(1):18802. doi:10.1038/s41598-020-75708-z

13. Zeng Y, Wei Z, Yu W, et al. Spatial transcriptomics prediction from histology jointly through Transformer and graph neural networks. Brief Bioinform. 2022;23(5):bbac297. doi:10.1093/bib/bbac297

14. Pang M, Su K, Li M. Leveraging information in spatial transcriptomics to predict super-resolution gene expression from histology images in tumors. Published online November 28, 2021:2021.11.28.470212. doi:10.1101/2021.11.28.470212

15. Levy JJ, Bobak CA, Nasir-Moin M, et al. Mixed Effects Machine Learning Models for Colon Cancer Metastasis Prediction using Spatially Localized Immuno-Oncology Markers. Pac Symp Biocomput Pac Symp Biocomput. 2022;27:175–186.

16. Svensson V, Teichmann SA, Stegle O. SpatialDE: identification of spatially variable genes. Nat Methods. 2018;15(5):343–346. doi:10.1038/nmeth.4636

17. Szegedy C, Vanhoucke V, Ioffe S, Shlens J, Wojna Z. Rethinking the Inception Architecture for Computer Vision. Published online December 11, 2015. doi:10.48550/arXiv.1512.00567

18. Dosovitskiy A, Beyer L, Kolesnikov A, et al. An Image is Worth 16×16 Words: Transformers for Image Recognition at Scale. In:; 2020. Accessed August 28, 2021. https://openreview.net/forum?id=YicbFdNTTy

19. Chen EY, Tan CM, Kou Y, et al. Enrichr: interactive and collaborative HTML5 gene list enrichment analysis tool. BMC Bioinformatics. 2013;14(1):128. doi:10.1186/1471-2105-14-128

20. McInnes L, Healy J, Saul N, Großberger L. UMAP: Uniform Manifold Approximation and Projection. J Open Source Softw. 2018;3(29):861. doi:10.21105/joss.00861

21. McInnes L, Healy J, Astels S. hdbscan: Hierarchical density based clustering. J Open Source Softw. 2017;2(11):205. doi:10.21105/joss.00205

22. Veen HJ van, Saul N, Eargle D, Mangham SW. Kepler Mapper: A flexible Python implementation of the Mapper algorithm. J Open Source Softw. 2019;4(42):1315. doi:10.21105/joss.01315

23. Tauzin G, Lupo U, Tunstall L, et al. giotto-tda: A Topological Data Analysis Toolkit for Machine Learning and Data Exploration. ArXiv200402551 Cs Math Stat. Published online April 6, 2020. Accessed July 23, 2020. http://arxiv.org/abs/2004.02551

24. Singh G, Mémoli F, Carlsson G. Topological Methods for the Analysis of High Dimensional Data Sets and 3D Object Recognition.:10.

25. Zhu J, Sun S, Zhou X. SPARK-X: non-parametric modeling enables scalable and robust detection of spatial expression patterns for large spatial transcriptomic studies. Genome Biol. 2021;22(1): 184. doi:10.1186/s13059-021-02404-0

26. Richards FM, McKee SA, Rajpar MH, et al. Germline E-cadherin Gene (CDH1) Mutations Predispose to Familial Gastric Cancer and Colorectal Cancer. Hum Mol Genet. 1999;8(4):607–610. doi:10.1093/hmg/8.4.607

27. Sawazaki S, Oshima T, Sakamaki K, et al. Clinical Significance of Tensin 4 Gene Expression in Patients with Gastric Cancer. In Vivo. 2017;31(6):1065–1071. doi:10.21873/invivo.11171

28. Pai SG, Carneiro BA, Mota JM, et al. Wnt/beta-catenin pathway: modulating anticancer immune response. J Hematol OncolJ Hematol Oncol. 2017;10(1):101. doi:10.1186/s13045-017-0471-6

29. Huang MS, Fu LH, Yan HC, et al. Proteomics and liquid biopsy characterization of human EMT-related metastasis in colorectal cancer. Front Oncol. 2022;12:790096. doi:10.3389/fonc.2022.790096

30. Li X, Li Z, Gu S, Zhao X. A pan-cancer analysis of collagen VI family on prognosis, tumor microenvironment, and its potential therapeutic effect. BMC Bioinformatics. 2022;23:390. doi:10.1186/s12859-022-04951-0

31. van Huizen NA, Coebergh van den Braak RRJ, Doukas M, Dekker LJM, IJzermans JNM, Luider TM. Up-regulation of collagen proteins in colorectal liver metastasis compared with normal liver tissue. J Biol Chem. 2019;294(1):281–289. doi:10.1074/jbc.RA118.005087

